# Novel Rhabdovirus and an almost complete drain fly transcriptome recovered from two independent contaminations of clinical samples

**DOI:** 10.1101/645325

**Authors:** Francisco Brito, Mosè Manni, Florian Laubscher, Manuel Schibler, Mary-Anne Hartley, Kristina Keitel, Tarsis Mlaganile, Valerie d’Acremont, Samuel Cordey, Laurent Kaiser, Evgeny M Zdobnov

## Abstract

Metagenomic approaches enable an open exploration of microbial communities without requiring a priori knowledge of a sample’s composition by shotgun sequencing the total RNA or DNA of the sample. Such an approach is valuable for exploratory diagnostics of novel pathogens in clinical practice. Yet, one may also identify surprising off-target findings. Here we report a mostly complete transcriptome from a drain fly (likely *Psychoda alternata*) as well as a novel Rhabdovirus-like virus recovered from two independent contaminations of RNA sequencing libraries from clinical samples of cerebral spinal fluid (CSF) and serum, out of a total of 724 libraries sequenced at the same laboratory during a 2-year time span. This drain fly genome shows a considerable divergence from previously sequenced insects, which may obscure common clinical metagenomic analyses not expecting such contaminations. The classification of these contaminant sequences allowed us to identify infected drain flies as the likely origin of the novel Rhabdovirus-like sequence, which could have been erroneously linked to human pathology, had they been ignored.

## Introduction

Metagenomic approaches allow us to comprehensively characterize the diversity of organisms in a sample without a priori knowledge of its content, and in a culture-independent manner. Thus it is rapidly becoming an indispensable tool for the characterization of novel and highly divergent organisms. Such an open approach also comes with accidental and/or unexpected findings. Several viruses have been identified in shotgun sequencing libraries, whose origin can be traced back to contaminants in reagents and/or library preparation errors. This issue becomes particularly important when handling patient libraries, since the identified organisms can be thought to be associated with disease etiology, potentially wasting time and resources in studies trying to link them. Of note, xenotropic murine leukemia virus was identified as a possible source of several diseases and health complications, only to be ruled out as a laboratory contaminant (1). Parvo-like hybrid viruses and kadipiro virus have also been identified as a contaminant of nucleic acid extraction spin columns from the QIAmp Viral RNA mini kit (Qiagen) (2,3). Other non-viral organisms have been erroneously identified as being part of the real sample, such as in the case of the microbiome of the placenta, which was shown to have been almost entirely comprised of background contaminants from the sequencing process (4,5). Bearing this in mind, a focus on the identification of unexpected contaminants is essential and takes precedence to other analyses, since it enables us to identify what is the actual metagenomic content of the sample, providing vital context for any downstream results.

Here we describe the analysis and identification of an accidental contamination by a drain fly and its associated virus in two sequencing libraries from clinical samples, highlighting the importance of performing in-depth genomic analyses in order to recognize unexpected, punctual contaminations, which can be missed even by setting up negative controls (6). Drain flies of the genus Psychoda (Psychodidae family), also known as moth flies, are ubiquitously found near water sources worldwide, infesting house drain pipes, sewage treatment plants, and even hospitals (7–9). Some species of Psychoda have also been associated to rare cases of human diseases, such as myiasis (10) and asthma (11), while other genera in the Psychodidae family have been found to be vectors of pathogenic viruses to both humans and other mammals, namely Rhabdoviruses (12,13). These are negative-sense single-stranded RNA viruses (ssRNA), which can also integrate into the genome of arthropods (14–17). In a clinical context, some Rhabdoviruses are also important human pathogens, causing diseases such as rabies and encephalitis (18,19). Indeed, novel rhabdoviruses have been already reported in clinical samples, both associated with disease (20), and in healthy individuals (21). Combining (meta)genomic and phylogenomics approaches, we assembled a mostly complete transcriptome of the drain fly *Psychoda alternata* from the two contaminated clinical samples, allowing us to identify the real source of the Rhabdovirus-like sequence which is highly divergent from any previously sequenced reference.

## Methods

### Sample collection, extraction and sequencing

Two clinical samples were analysed: one cerebrospinal fluid (CSF) specimen collected in 2014 at the University Hospitals of Geneva, Switzerland, from a 59-year-old patient hospitalized for meningitis of unknown origin, and one serum collected in 2015 in Dar es Salaam, United Republic of Tanzania, from a 22.5-month old child presenting a fever of 40.1°C and reporting abdominal pain without other gastrointestinal symptoms (malaria rapid test was positive and the fever resolved within 2 days of antimalarial therapy). RNA was extracted from the cerebral spinal fluid (CSF) and the serum samples as previously described (i.e. centrifugation, DNAse treatment and ribonucleic acid extraction by TRIzol) (22) in 2015 and 2017, respectively. RNA libraries were prepared using the TruSeq total RNA preparation protocol (Illumina, San Diego, US). Libraries were run on a Hiseq2500 platform (Illumina) using the 2×100-nucleotide read length protocol.

### Read filtering and analysis

Illumina adapters were removed and read quality was assessed using Trimmomatic v0.33 (23). Human data was removed by mapping reads against the human genome (hg38) using SNAP v1.0beta.23. (24). Reads were assessed for complexity with tagdust2 v2.33 (25), then mapped against the uniVec database in order to exclude possible reagent contaminants, and against RefSeq’s ribosomal RNA databases for bacteria and fungi (28S, 23S, 18S, 16S, and 5S). The filtered reads were assembled using MEGAHIT v1.1.12 (26). Recovered contigs were aligned against NR (as of Feb 2019) using DIAMOND v0.9.13.114 (27) and binned using MEGAN v6.15.0 (28). Recovered viral contigs were translated using the Expasy translate tool (29) and the obtained amino acid sequences were aligned against NR using PSI-Blast, up to 5 iterations (30). Coverage of viral contigs was calculated by mapping back the reads to the contigs, using SNAP.

### Insect identification

To identify the arthropod species source of the contamination, mitochondrial cytochrome c oxidase subunit I *(COI)* genes were identified with BLAST searches against the *COI* genes of several arthropods, and significant hits were searched against the BOLD System (Barcode Of Life Data System) (31). Contigs classified by MEGAN as arthropod, or higher than arthropod but whose top hits were still arthropods, were analysed with BUSCO v3 (32) to estimate the completeness of the assembled transcriptome using the Arthropoda, Insecta, and Diptera datasets. To obtain markers for building a phylogenetic tree, 121 single-copy genes were extracted from *Psychoda alternata* contigs and the genomes of 19 other arthropods (17 Diptera, 1 Lepidoptera and 1 Hymenoptera) using the scripts from BUSCO available at https://gitlab.com/ezlab/busco_usecases/tree/master/phylogenomics. The arthropod genomes used were *Acromyrmex echinatior, Bombyx mori, Culex quinquefasciatus, Aedes aegypti, Aedes albopictus, Anopheles gambiae, Anopheles minimus, Anopheles culicifacies, Polypedilum vanderplanki, Polypedilum nubifer, Mayetiola destructor, Drosophila grimshawi, Drosophila ananassae*, *Drosophila erecta, Musca domestica, Glossina palpalis*, *Glossina austeni, Rhagoletis zephyria* and *Phlebotomus papatasi*. Briefly, single-copy genes from each species were aligned individually with MAFFT v7 (33) and each multiple sequence alignment (MSA) was subsequently trimmed with Trimal v1.4 using the “-auto” option. The MSAs were then concatenated in a supermatrix of 76480 aa, which was used to infer a maximum likelihood tree with RaxML v8.2.11 (34) using the PROTGAMMAJTT model. We estimated 100 bootstrap replicates and bootstrap values were drawn on the best ML tree using the rapid bootstrapping RAxML option. The tree was visualised with EvolView (35). Proteins were predicted by translating the recovered contigs in all 6 frames and recovering the longest ORF for each. These were then mapped against OrthoDB (36) using the website’s analysis feature, and comparative charts of the number of orthologous genes recovered against a set of 8 representative dipteran species were generated.

### Virus analysis

Virus phylogeny was made by performing an MSA at the amino acid level with 26 known rhabdovirus L proteins (also known as RNA dependant RNA polymerase) and the two predicted L proteins – one from each – recovered rhabdovirus sequence. The ML tree was inferred using IQ-TREE v.1.5.5 (37), with a LG+F+I+G4 substitution model, and using 1000 bootstrap replicates. Variant calling between the two novel Rhabdovirus sequences was performed by mapping back the MG2017 reads against the MG2015 viral sequence and calling variants with Lofreq2 v2.1.2 (38). To confirm the absence of the viruses in the human metagenomic samples, a real-time RT-PCR, specific for the detection of the two novel rhabdovirus-like sequences, was designed (forward primer 5’-TGCCCCCCTGGTTACCA-3’, reverse primer 5’-CCGGCTGCATCAGGATCT-3’, and probe 5-FAM-TGTTCCCATCCGCATAT-MGB NFQ-3’). RNA were extracted from the two initial positive samples using the NucliSENS easyMAG (bioMérieux, Geneva, Switzerland) nucleic acid kit, and then tested by real-time RT-PCR using the one-step QuantiTect Probe RT-PCR Kit (Qiagen, Hombrechtikon, Switzerland) in a StepOne Plus instrument (Applied Biosystems, Rotkreuz, Switzerland) under the following cycling conditions: 50 °C for 30 min, 95°C for 15 min, 45 cycles of 15 s at 94 °C and 1 min at 55°C. In parallel, cDNA was randomly synthesized (random hexamers) using the reverse transcriptase SuperScript II (Invitrogen, Carlsbad, CA, USA) and then tested by PCR (forward primer 5’-CAGGATCTTATATGCGGATGGGAACAGT-3’ and reverse primer 5’-CTCTTAGGAAAGAAGGCCTTCATGGACCT-3’; expected fragment size = 333).

## Results and Discussion

The two contaminated RNA metagenomic libraries are referred to as MG2015 and MG2017, describing the years of sequencing of the samples (4 March 2015 and 15 November 2017). No other similar contamination events were observed across the 724 library preparations made between the contaminated samples’ preparation dates, in the same laboratory. After quality filtering and removal of human data, both libraries show a high amount of non-human reads: 88% of the total initial reads in library MG2017 and 46% in library MG2015 (Table 1). Contig binning classified 9468 contigs in library MG2017 and 1402 contigs in library MG2015 as arthropoda-like contigs, which comprise 72% and 8% of total filtered reads, respectively. Remaining reads consist of bacterial data, viruses and divergent human data not caught by the initial filter. Mapping the MG2015 reads to the MG2017 arthropod contigs significantly increased the number of mapped reads (49%), suggesting low transcriptome coverage in MG2015. In order to identify which species of arthropod was present on each library, we recovered – from both assemblies – the full sequences of the *COI* and compared them against all barcode records on the Barcode Of Life Database (BOLD). Both *COI* sequences display a 100% nucleotide identity to *COI* of the drain fly *Psychoda alternata* (Diptera: Psychodidiae) (Sup. Figure 1). We further assessed the recovered drain fly transcriptomes by checking their completeness in terms of expected gene content, using BUSCO. The transcriptome from library MG2017 harbours 84.8%, 79.4% and 54.3% of the single-copy genes expected to be present in arthropods, insects and dipterans, respectively (Sup. Figure 2). A substantially lower number of single-copy genes were present in library MG2015 (2.8%, 2.8% and 2.4% for arthropods, insects and dipterans, respectively) reflecting the lower number of assembled contigs. Merging both libraries together and re-assembling marginally improves the BUSCO scores to 85.3%, 79.9% and 55% (Arthropoda, Insecta and Diptera). The phylogenetic tree inferred using 121 single-copy genes from library MG2017 and 20 other insect genomes placed the contaminant insect species within the clade of moth flies Psychoda (Figure 1), corroborating the result obtained by comparing the barcoding sequence of *COI.* We expanded on this analysis by taking all predicted proteins from the suspected *P. alternata* transcriptome and mapping them to proteins from dipteran species in OrthoDB. The comparative chart built on OrthoDB website (Figure 2) shows that more than 6000 of *P. alternata*’s predicted proteins have a corresponding ortholog in at least another of the 8 selected dipteran species. Of these, c.a. 1000 are likely to be present in single-copy in all species, c.a. 400 are single-copy in almost all species and the remaining are present in multiple copies across the selected species.

**Figure 1.**
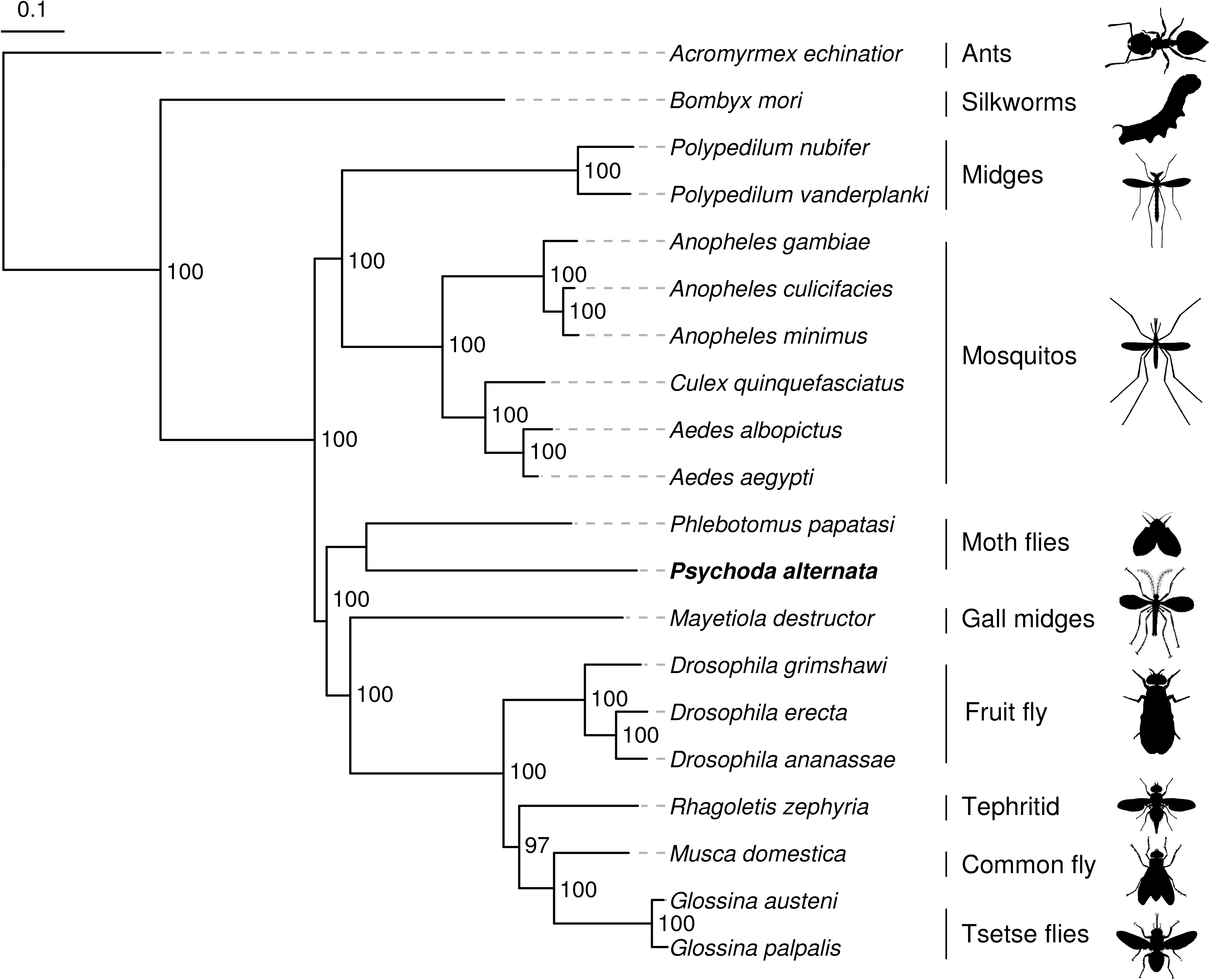
Maximum likelihood phylogeny of the recovered *Psychoda alternata* contigs and the genomes of 19 other arthropods (17 Diptera, 1 Lepidoptera and 1 Hymenoptera) based on aligned protein sequences of 121 single-copy orthologs. Branch lengths represent substitutions per site. Values on nodes indicate bootstrap support.

**Figure 2.**
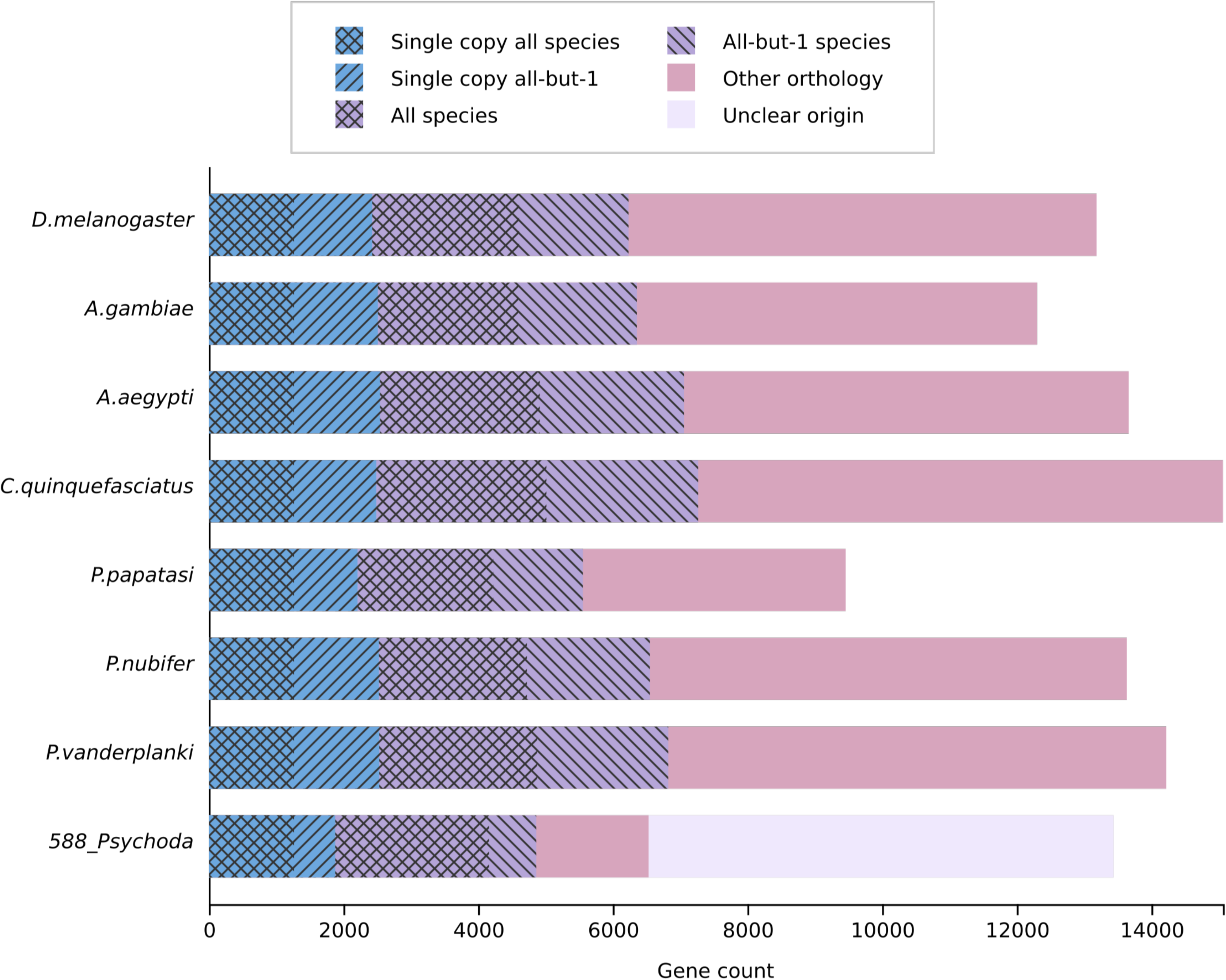
OrthoDB comparative chart of the number of predicted orthologous genes from the assembled *P. alternata* transcriptome and the gene sets of 8 representative dipteran species.

**Table 1.** Read statistics, number of arthropod contigs and Rhabdovirus-like contigs on libraries MG2015 and MG2017.

|  | MG2017 |  | MG2015 |  |
| --- | --- | --- | --- | --- |
|  | No. | % | No. | % |
| Raw reads | 161,759,116 | 100.00 | 113,183,610 | 100.00 |
| Non-human reads | 142,527,300 | 88.11 | 51,975,118 | 45.92 |
| rRNA filtered, non-human reads | 130,489,704 | 80.67 | 47,818,244 | 42.25 |
| Arthropod contigs | 9,468 | - | 1,402 | - |
| Reads mapping to MG2017 arthropod contigs | 93,915,071 | 58.06 | 23,519,926 | 20.78 |
| Reads mapping to MG2015 arthropod contigs | 6,346,874 | 3.92 | 3,952,716 | 3.49 |
| Rhabdovirus-like virus contigs | 1 | - | 1 | - |
| Reads mapping to rhabdovirus-like virus contigs | 80,973 | 0.05 | 38,284 | 0.03 |

The presence of Psychodidae in Europe and Switzerland (where the samples were processed) is well documented, with several species being described in western Switzerland alone (39). Its ubiquity and its pervasiveness in drain pipes make it a potential contamination hazard in laboratories and hospitals, the latter where they have been reported infesting an operating room (9), though no previous reports of drain fly contamination have been made in other experiments.

We also identified a Rhabdovirus-like sequence in both libraries, with a sequence similarity of 99.4% (GC content: 48.5%) between them (Figure 3). The MG2015 sequence has a length of 11958 nt and an average read depth of 628, whereas MG2017 has a length of 11941 nt and an average depth of 655 (Sup. Figure 3). These values are within the expected length for Rhabdoviruses, which ranges from 11000 to 16000 nt (16). Both sequences are comprised of 5 hypothetical proteins, 4 of them having similarities to the canonical rhabdovirus proteins (N, MP, G, L) (12). PSI-Blast results show an overall sequence similarity of 22% at the amino acid level (N: 21% Kotonkan virus YP_006202618.1, MP: 14.5% Nishimuro ledantevirus YP_009505473.1, G: 19% Vesicular Stomatitis Indiana Virus ACK77584.1, and L: 33% Vesicular Stomatitis New Jersey Virus AUI41073.1, each covering 97-99% of the queried sequences). Although a fifth protein is located in the same region where a fifth canonical protein (protein P) is commonly found in other rhabdovirus genomes, no sequence similarity was found to any known protein. At the nucleotide level, the closest sequence is a *Pararge aegeria* (butterfly) rhabdovirus (KR822826.1), covering only 5% of the total sequence. Between both newly-discovered rhabdovirus-like sequences, we find 42 SNPs with an allelic frequency of 1, distributed along the genome (Sup. Table 2), causing a total of 5 amino acid changes across all predicted proteins. The SNPs observed do not change the length of the predicted proteins, and are likely to derive from natural variation of the same viral species, across two years of infecting different flies.

**Figure 3.**
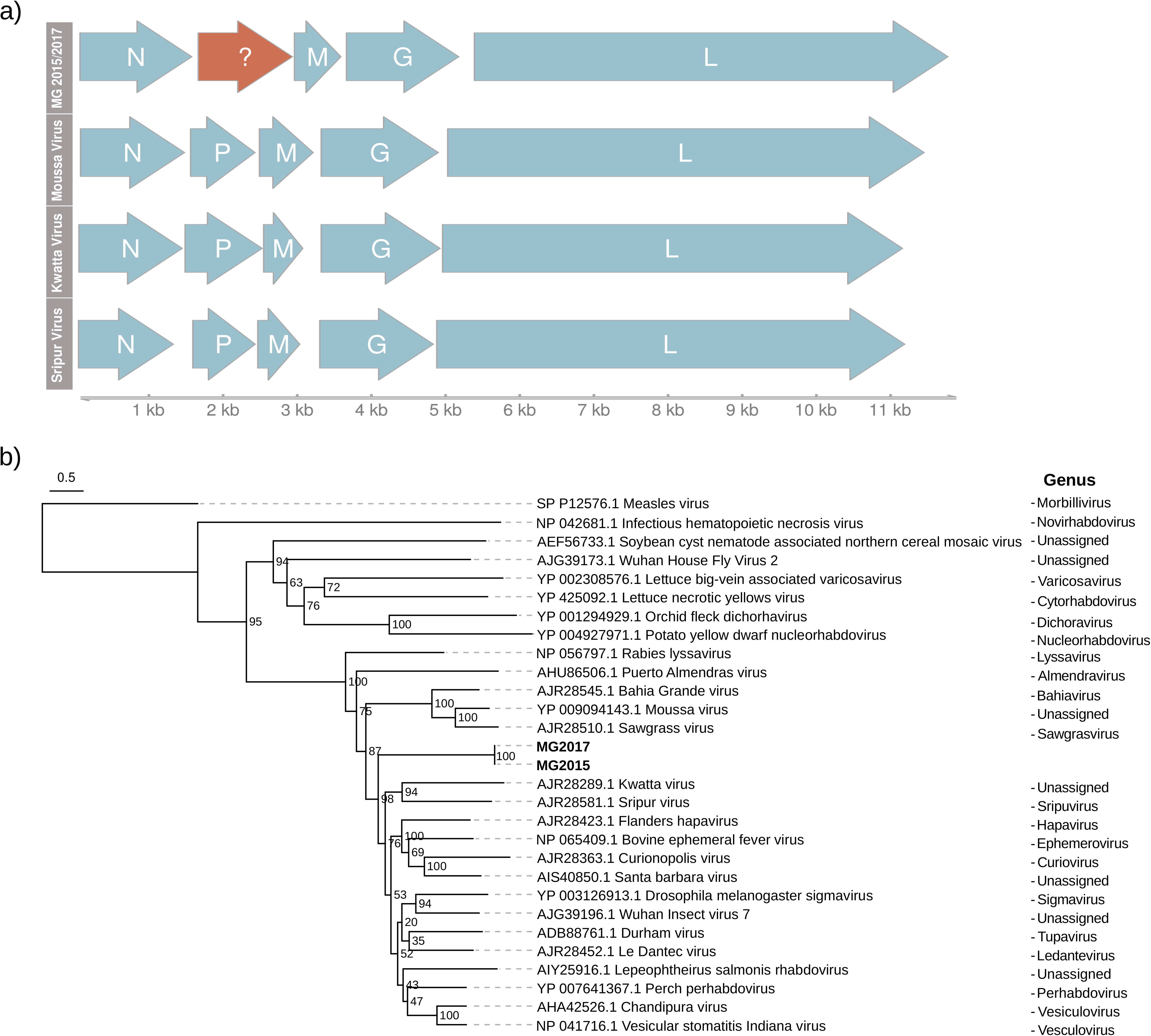
a) Schematic representation comparing the ORFs of the recovered Rhabdovirus-like sequence with the five canonical proteins present in three close known Rhabdovirus genomes – Kwatta virus (KM204985.1), Sripur virus (NC_034542.1) and Moussa virus (NC_025359.1)- b) Maximum likelihood phylogeny of 26 L-protein sequences of previously reported rhabdoviruses and the predicted L-protein sequence of the newly recovered rhabdovirus-like virus. Branch lengths represent substitutions per site. Values on nodes indicate bootstrap support.

The presence of this virus in the samples was not confirmed by qPCR, suggesting the Rhabdovirus is associated to the Psychoda contaminant, either by infection, or integration into the genome as described in Geisler & Jarvis (17). Since rhabdoviruses can infect a broad range of organisms, other than humans (13–16), the detection of rhabdoviruses in human samples requires a thorough inspection to avoid reporting any false positive discoveries, which would erroneously associate species of rhabdoviruses to human health. We also analysed the DNA libraries generated from the same original samples (not shown/unpublished). Both are negative for insect and virus, pointing towards the contamination occurring specifically during RNA library preparation.

Using an open approach, we identified a complex contamination case: a novel virus resembling species linked to human disease (Rhabdoviruses) infecting a never before sequenced insect (*P. alternata*), which in turn was contaminating clinical samples. This required us to use a phylogenomics approach in order to correctly classify the insect contaminant before being able to accurately determine the origin of the virus infection. While it only affected 2 out of 724 library preparations (0.27%), due to the ubiquity of the drain fly, it is possible that this unusual source of contamination could be observed in other laboratories. These results show us the potential impact of contaminants on the interpretation of clinical samples, and how they could bias how we classify organisms as possibly pathogenic, ultimately shifting the focus to signals that are irrelevant to the actual patients, leading to wrong conclusions, and potentially to misdiagnosis.

## Acknowledgements

We thank Mylène Docquier and Brice Petit from the iGE3 Genomics Platform, University of Geneva, Switzerland. The study was supported by the Bill and Melinda Gates Foundation funding OPP1163434 to VDA, the Swiss National Science Foundation funding 32003B_146993 to LK and 31003A_166483 to EZ, and the IGE3 Award to FB.

## Authors’ contributions

MS, MAH, KK, TM, VDA, LK, SC collected the clinical samples, performed the HTS sample preparations, the PCR analysis, funded the sequencing of these libraries and reviewed the manuscript. FB and FL co-discovered the Rhabdovirus and analysed the metagenomic data. FB, MM, and EZ further analysed and interpreted the data, and FB, MM, SC and EZ wrote the manuscript.

## Availability of data and materials

Recovered rhabdovirus-like sequences, COI and arthropod contigs are available at: http://cegg.unige.ch/contamination_metagenomics

---

**Supplementary Figure 1 –** Phylogenetic identification of the recovered COI genes in a) MG2015, b) MG2017 against the BOLD database. Figure generated using the website’s Tree Identification feature.

**Supplementary Figure 2 –** Busco assessment results using the Arthropoda single-copy gene set for the MG2015 and MG2017 libraries, and using the Diptera gene set for MG2015, MG2017, and the closest annotated dipteran species *(Phlebotomus papatasi).*

**Supplementary Figure 3 –** Read depth and coverage of the Rhabdovirus-like sequences from a) MG2015 and b) MG2017 libraries. Below each coverage plot, the location of each protein is represented. N – Nucleocapsid, G – Glycoprotein, ? – No known similarities to annotated proteins, M – Matrix protein, L – RNA dependent RNA polymerase.

**Supplementary Table 1 –** Table of variants found in the MG2017 rhabdovirus-like virus, compared against the MG2015 rhabdovirus-like virus.

## Notes

http://cegg.unige.ch/contamination_metagenomics

