## Supplemental Figures 1-3 and Supplemental Table 1 for "Novel Rhabdovirus and an almost complete drain fly transcriptome recovered from two independent contaminations of clinical samples"

Supplementary Figure 1

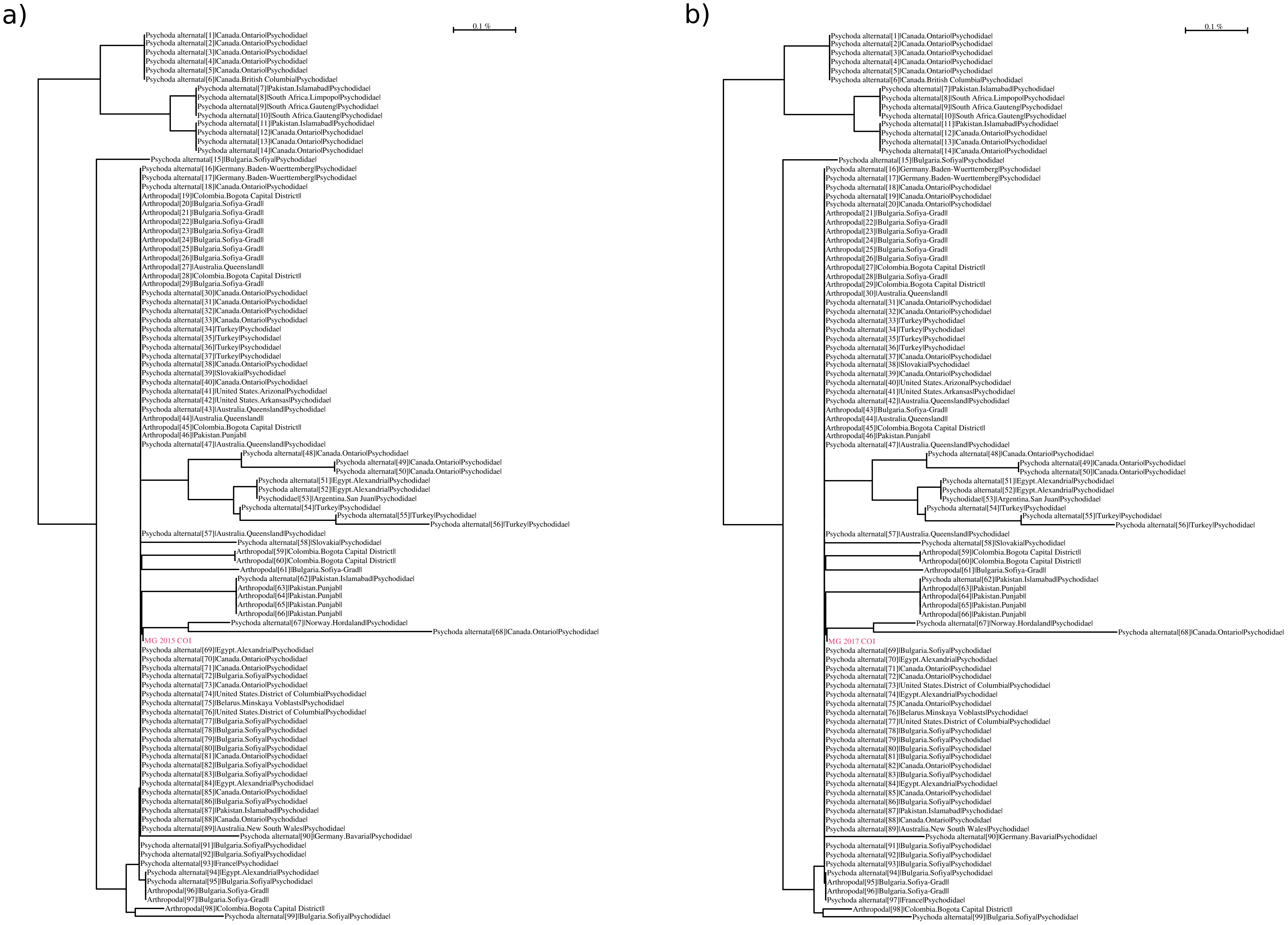

Supplementary Figure 2

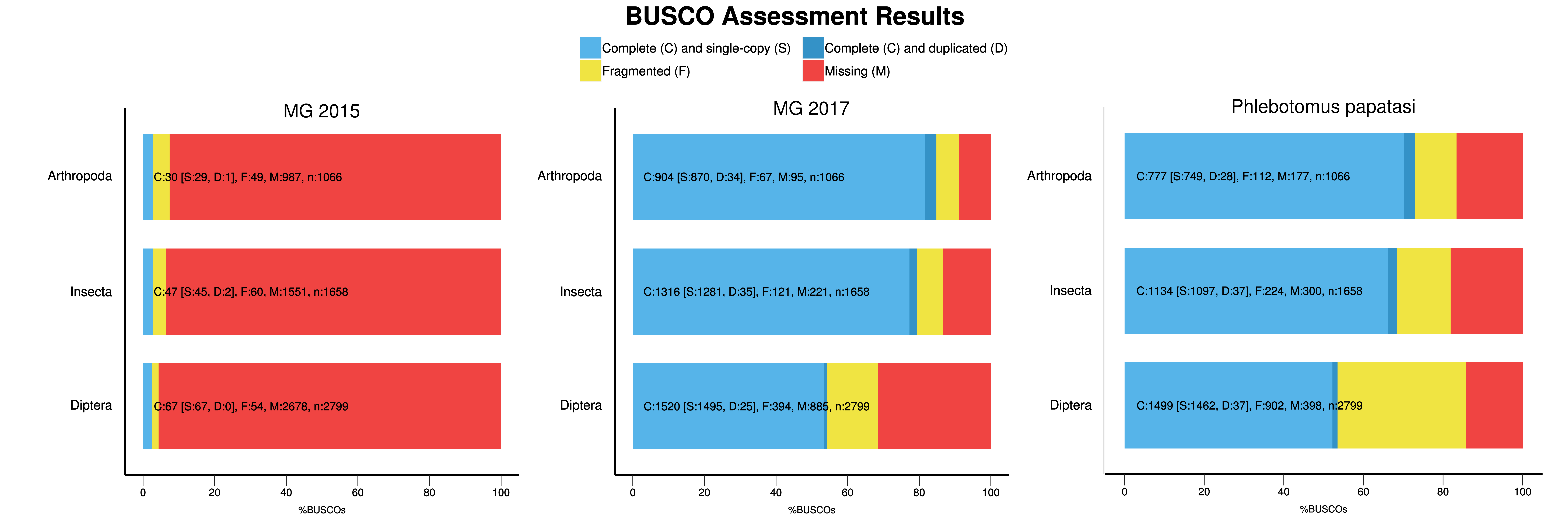

Supplementary Figure 3

a)

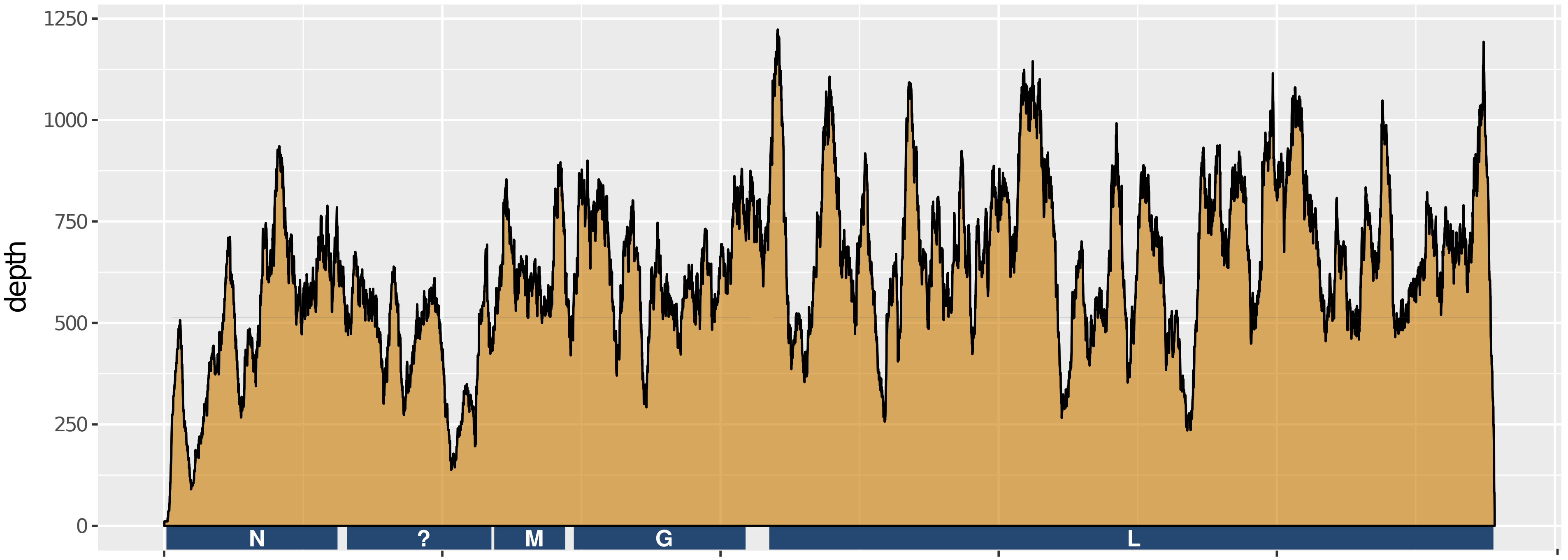


MG2015 | genome length: 11958 bp | average depth: 320 | total covered bp: 11958

b)

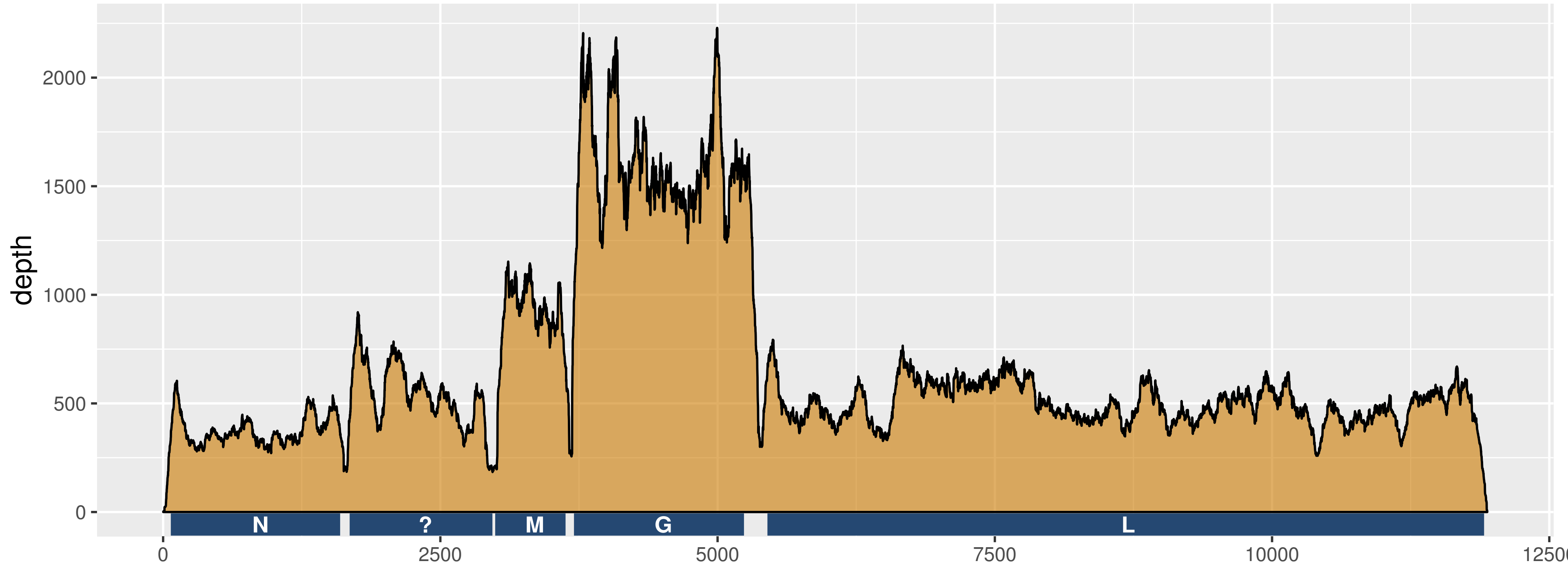


MG2017 | genome length: 11941 bp | average depth: 655 | total covered bp: 11941

Supplementary Table 1

##fileformat=VCFv4.0
##fileDate=20190228
##source=lofreq call --call-indels -f MG2015.fa -o Dindel.MG2017.vcf Dindel.MG2017.bam
##reference=MG2015.fa
##INFO=<ID=DP,Number=1,Type=Integer,Description="Raw Depth">
##INFO=<ID=AF,Number=1,Type=Float,Description="Allele Frequency">
##INFO=<ID=SB,Number=1,Type=Integer,Description="Phred-scaled strand bias at this position">
##INFO=<ID=DP4,Number=4,Type=Integer,Description="Counts for ref-forward bases, ref-reverse, alt-forward and alt-reverse bases">
##INFO=<ID=INDEL,Number=0,Type=Flag,Description="Indicates that the variant is an INDEL.">
##INFO=<ID=CONSVAR,Number=0,Type=Flag,Description="Indicates that the variant is a consensus variant (as opposed to a low frequency variant).">
##INFO=<ID=HRUN,Number=1,Type=Integer,Description="Homopolymer length to the right of report indel position">
##FILTER=<ID=min_dp_10,Description="Minimum Coverage 10">
##FILTER=<ID=sb_fdr,Description="Strand-Bias Multiple Testing Correction: fdr corr. pvalue > 0.001000">
##FILTER=<ID=min_snvqual_60,Description="Minimum SNV Quality (Phred) 60">
##FILTER=<ID=min_indelqual_46,Description="Minimum Indel Quality (Phred) 46">
#CHROMPOSIDREFALTQUALFILTERINFO
MG2015_len=11958151.AC6157PASSDP=184;AF=1.000000;SB=0;DP4=0,0,106,78
MG2015_len=11958308.CT7405PASSDP=211;AF=0.995261;SB=0;DP4=0,0,108,102
MG2015_len=11958319.GA7646PASSDP=220;AF=1.000000;SB=0;DP4=0,0,106,114
MG2015_len=11958471.CT74PASSDP=211;AF=0.023697;SB=0;DP4=126,80,3,2
MG2015_len=11958518.CT2806PASSDP=203;AF=0.512315;SB=7;DP4=52,47,44,60
MG2015_len=11958550.CT6706PASSDP=196;AF=0.989796;SB=0;DP4=0,0,89,105
MG2015_len=119581180.TC5156PASSDP=144;AF=1.000000;SB=0;DP4=0,0,78,66
MG2015_len=119581639.TC6943PASSDP=191;AF=1.000000;SB=0;DP4=0,0,112,79
MG2015_len=119581732.CT82PASSDP=183;AF=0.027322;SB=0;DP4=101,77,3,2
MG2015_len=119582001.CT102PASSDP=236;AF=0.029661;SB=5;DP4=106,123,5,2
MG2015_len=119582125.GT6788PASSDP=192;AF=0.994792;SB=0;DP4=0,0,103,88
MG2015_len=119582345.CTC69PASSDP=185;AF=0.016216;SB=5;DP4=81,101,0,3;INDEL;HRUN=1
MG2015_len=119582530.TC5720PASSDP=156;AF=1.000000;SB=0;DP4=0,0,74,82
MG2015_len=119582557.GA5365PASSDP=154;AF=1.000000;SB=0;DP4=0,0,83,71
MG2015_len=119582872.CT5200PASSDP=145;AF=0.986207;SB=0;DP4=0,0,61,82
MG2015_len=119583295.GA5116PASSDP=148;AF=0.993243;SB=0;DP4=0,0,73,74
MG2015_len=119583606.CT4892PASSDP=141;AF=1.000000;SB=0;DP4=0,0,61,80
MG2015_len=119583766.CT5738PASSDP=162;AF=0.993827;SB=0;DP4=0,0,77,84

MG2015_len=119583847.CT6005PASSDP=170;AF=0.994118;SB=3;DP4=0,1,90,79
MG2015_len=119584147.CT8386PASSDP=242;AF=1.000000;SB=0;DP4=0,0,125,117
MG2015_len=119584183.AG8982PASSDP=249;AF=1.000000;SB=0;DP4=0,0,145,104
MG2015_len=119584762.TGT68PASSDP=190;AF=0.015789;SB=0;DP4=97,95,2,1;INDEL;HRUN=1
MG2015_len=119585011.TC7678PASSDP=216;AF=0.995370;SB=0;DP4=0,1,93,122
MG2015_len=119585164.GA7849PASSDP=219;AF=1.000000;SB=0;DP4=0,0,96,123
MG2015_len=119585203.AC471PASSDP=229;AF=0.109170;SB=5;DP4=100,104,9,16
MG2015_len=119585275.TC7639PASSDP=217;AF=1.000000;SB=0;DP4=0,0,99,118
MG2015_len=119585644.CT7177PASSDP=203;AF=0.995074;SB=3;DP4=0,1,111,91
MG2015_len=119585892.GT5162PASSDP=144;AF=1.000000;SB=0;DP4=0,0,72,72
MG2015_len=119585960.GAG47PASSDP=142;AF=0.014085;SB=3;DP4=70,71,2,0;INDEL;HRUN=1
MG2015_len=119586018.AG79PASSDP=169;AF=0.029586;SB=11;DP4=89,75,5,0
MG2015_len=119586302.GC5945PASSDP=170;AF=1.000000;SB=0;DP4=0,0,68,102
MG2015_len=119586334.GA6495PASSDP=182;AF=1.000000;SB=0;DP4=0,0,108,74
MG2015_len=119586460.GA7001PASSDP=193;AF=1.000000;SB=0;DP4=0,0,69,124
MG2015_len=119586622.AG64PASSDP=278;AF=0.025180;SB=2;DP4=233,37,7,0
MG2015_len=119586810.CA23330PASSDP=683;AF=0.995608;SB=3;DP4=1,0,331,349
MG2015_len=119586883.AG64PASSDP=661;AF=0.015129;SB=27;DP4=330,320,10,0
MG2015_len=119587221.AC20751PASSDP=603;AF=0.993366;SB=3;DP4=1,0,281,318
MG2015_len=119587752.AG20908PASSDP=608;AF=0.995066;SB=0;DP4=1,1,232,373
MG2015_len=119587997.AC22383PASSDP=620;AF=0.996774;SB=0;DP4=0,0,228,390
MG2015_len=119588960.GA1854PASSDP=51;AF=1.000000;SB=0;DP4=0,0,22,29
MG2015_len=119589294.TC6450PASSDP=183;AF=1.000000;SB=0;DP4=0,0,126,57
MG2015_len=119589567.GA6404PASSDP=174;AF=1.000000;SB=0;DP4=0,0,87,87
MG2015_len=119589701.CT6548PASSDP=191;AF=1.000000;SB=0;DP4=0,0,106,85
MG2015_len=119589738.GA6860PASSDP=197;AF=1.000000;SB=0;DP4=0,0,111,86
MG2015_len=119589841.CA8846PASSDP=262;AF=1.000000;SB=0;DP4=0,0,132,130
MG2015_len=1195810400.GA5136PASSDP=146;AF=0.993151;SB=0;DP4=1,0,104,41
MG2015_len=1195810557.CT67PASSDP=136;AF=0.029412;SB=0;DP4=63,68,2,2
MG2015_len=1195810735.TCT48PASSDP=119;AF=0.016807;SB=3;DP4=55,63,0,2;INDEL;HRUN=1
MG2015_len=1195810805.AGA48PASSDP=123;AF=0.016260;SB=7;DP4=69,54,0,2;INDEL;HRUN=1
MG2015_len=1195810877.TC3917PASSDP=114;AF=1.000000;SB=0;DP4=0,0,53,61
MG2015_len=1195811214.AG80PASSDP=128;AF=0.039062;SB=7;DP4=51,72,4,1
MG2015_len=1195811321.GA4840PASSDP=143;AF=1.000000;SB=0;DP4=0,0,70,73

MG2015_len=1195811432.CT4108PASSDP=116;AF=1.000000;SB=0;DP4=0,0,70,46
MG2015_len=1195811509.AG4634PASSDP=130;AF=1.000000;SB=0;DP4=0,0,61,69
MG2015_len=1195811537.GA4086PASSDP=118;AF=0.991525;SB=3;DP4=1,0,52,65
MG2015_len=1195811584.GA2984PASSDP=89;AF=1.000000;SB=0;DP4=0,0,41,48
